# Pupillary aperture is a potential biomarker of movement preparation

**DOI:** 10.1101/2021.01.30.428981

**Authors:** Pragya Pandey, Supriya Ray

## Abstract

In response to variable light intensity, the pupils reflexively constrict or dilate to maintain a uniform retinal illumination. The pupillary light reflex (PLR) pathway receives projections from two important areas in primates’ brain that plan rapid saccadic eye-movement – frontal eye field (FEF) and superior colliculus (SC). The speed with which neurons in these areas increase firing rate to a threshold determines latency of a saccade. Micro-stimulation of FEF/SC neurons below this threshold modulates the magnitude of PLR. Nonetheless, how the saccade latency and pupil dynamics are related remains unknown. Our study shows that the appearance of a bright stimulus evokes pupil constriction at higher rate when the latency of impending saccade to the stimulus is shorter. This inverse relationship between the rate of pupil constriction and the saccade latency is robust irrespective of the reward outcome. In a homeomorphic biomechanical model of pupil, we have projected build-up signal similar to FEF and SC activity to the parasympathetic and sympathetic divisions of the PLR pathway, respectively. Model simulation mimics the observed data to indicate that the FEF and SC activity for eye movement modulates autonomic input to the pupillary muscle plant. A striking similarity between the dynamics of pupil constriction and stochastic rise in neural activity for saccade elicitation suggests that PLR is a potential proxy of movement preparation, and not mere an indicator of attentional orientation. Our study suggests a mechanism of how the retinal luminosity is timely regulated to aid perception by minimizing visual transients due to gaze orientation.

## INTRODUCTION

The luminosity of the visual space around us is heterogeneous, therefore the foveola in the retina may experience either overwhelming or impoverished visual transient after every cycle of rapid saccadic eye movement and gaze fixation. Saccade brings the object of interest on the fovea for visual processing in high resolution in larger spatial scale, whereas fixational microsaccades process the foveated object in finer scale for thorough inspection, which is highly sensitive to the contrast of the object (Boi et al., 2017; Intoy and Rucci, 2020). Our brain eliminates retinal blurring during saccade and maintains spatial stability after saccade by predictively suppressing visual acuity (Zuber and Stark, 1966; Ross et al., 2001; Thiele et al., 2002) and remapping the visual world to future receptive field of neurons (Colby and Goldberg, 1992; Bays and Husain, 2007; Melcher and Colby, 2008), respectively. In addition, visual adaptation by means of a timely adjustment of pupillary aperture before saccade elicitation may reduce the instability in retinal illumination induced by gaze shift and fixation cycle.

Before the elicitation of saccade, pupil indeed dilates if gaze orients towards dark, and constricts if gaze orients towards light. When an auditory cue instructed participants to orient gaze towards a bright or dark side on the left or the right side of the display, the pupil began to respond to the brightness of the cued side with near zero latency (i.e. while the eyes were in motion), suggesting simultaneous preparation for saccade and pupillary response (Mathôt et al., 2015). In gap task, wherein the central fixation spot disappears before onset of the peripheral target, larger pupil size accompanies with shorter saccade latency (Wang et al., 2015), and pupil dilation in a trial relative to mean dilation increases with decreasing saccade latency (Jainta et al., 2011). These findings suggest that the pupillary light response (PLR) and preparation for eye movement are linked.

Covert orientation of attention at the cued location without orienting the gaze to a bright (or dark) patch can also induce pupil constriction (or dilation) (Binda et al., 2013a; Mathôt et al., 2013; Binda and Murray, 2015). Pupil dynamics are similar when saccade orientation is guided by the memorized representation of the stimulus compared to when guided by the stimulus itself (Wang et al., 2018). These findings indicate that the covert shift of attention preceding eye movement, either obligatory (endogenous) or cued (exogenous), can trigger pupillary response. Covert pre-saccadic obligatory attentional shift occurs within a couple of hundreds of milliseconds from the beginning of saccade planning (Kröse and Julesz, 1989; Müller and Rabbitt, 1989; Nakayama and Mackeben, 1989; Carlson et al., 2006; Castet et al., 2006), if and only if gaze orientation is inevitable (Born et al., 2014). Attention is suppressed at the location where directing saccade is prohibited (Dhawan et al., 2013).

In primates’ brain, the frontal eye field (FEF) and the superior colliculus (SC) that majorly contribute to saccade planning and pre-saccadic covert orientation of attention, also send projections to the PLR pathway (Wang and Munoz, 2015). Activity of specific types of neurons in FEF and SC increase to a threshold to trigger saccade. The rate of increase in firing rate of these neurons determines saccade latency (Hanes and Schall, 1996; Dorris et al., 1997). Visuo-movement neurons in the FEF/SC respond after visual stimulus onset and subsequently before saccade elicitation. These neurons in the FEF exhibit movement related modulation in activity if saccade is inevitable (Ray et al., 2009), which to some extent supports behavioural finding that attention covertly shifts to the saccade end point if saccade is certain (Born et al., 2014). Previous micro-stimulation studies on monkeys showed that the FEF/SC activity influences pupil size. Sub-threshold electrical micro-stimulation of the FEF that did not elicit saccadic eye movement could increase pupil size (Lehmann and Corneil, 2016). However, subsequent study did not find the main effect of sub-saccadic micro-stimulation of FEF on pupil size. Rather weak micro-stimulation modulated the magnitude of pupil constriction evoked by a bright peripheral ‘probe’ stimulus when presented briefly inside the response field of the stimulated neuron (Ebitz and Moore, 2017). Similar sub-saccadic micro-stimulation of the intermediate layer in the subcortical superior colliculus (SCi) dilated the pupils. The magnitude of evoked pupil dilation was larger for dim background that bright background. In contrast, micro-stimulation of the superficial layer (SCs) containing visually responsive neurons did not affect the pupil size (Wang et al., 2012). Note that the SCi, where saccade related neurons are found in abundance, receives input from both the FEF and SCs, and projects to brainstem to generate saccade (White and Munoz, 2011). The above findings together suggest that PLR, saccadic reaction time (SRT), and the FEF/SC activity pertaining to saccade planning are linked. However, whether the neural dynamics of saccade planning and PLR are related remains unknown.

We hypothesized that pupillary aperture changes at faster rate for shorter latency saccade, and vice versa, for a timely pre-saccadic adjustment of the influx of light to reduce abrupt change in luminosity on the retinae after saccade. We examined whether the dynamics of change in pupil size and SRT are related on trial-by-trial basis. We distinguished influence of saccade preparation from mere shift of attention on pupillary response by contrasting pupil dynamics when saccades were executed against when planned saccades were cancelled. Our data suggest a robust inverse relationship between the rate of pupil constriction and saccade latency. We further sought to understand how the oculomotor system that plan a saccadic eye movement and the autonomic nervous system that drives iris muscles interplay to realize this relationship. To this end, we extended a homeomorphic biomechanical model of pupillary muscle plant (Usui and Hirata, 1995) by projecting rise-to-threshold activity of the FEF and SC to the parasympathetic and sympathetic division of the PLR pathway, respectively. Simulation of our neuromechanical model of pupil mimicked the dynamics of PLR and accounted for our empirical findings, indicating that the pupil size has potential to be a marker of neural activity pertaining to saccade planning.

## MATERIALS AND METHODS

### Participants

A total of 20 healthy humans (13 males) with correct or corrected to normal vision performed a choice countermanding task. The average (± *SD*) age of the participants was 21 (± 2.49) years. All participants gave their informed consent in writing to take part in this study in advance. They were naïve in the behavioral task and unaware of the scientific goal of the experiment. This study was in compliance with the Declaration of Helsinki (World Medical Association, 2008), and was approved by the guidelines of the Institutional Ethics Review Board of University of Allahabad for biomedical research on human participants. Participants received instructions to perform the task both verbally and in writing either in English or their native language.

### Apparatus

Participants sat on a chair comfortably and put their chin on a custom-made chin-forehead resting apparatus to minimize head movement. The stimuli were displayed on a 19 inch LCD monitor (resolution: 640 × 480, refresh rate: 60 Hz, Aspect ratio: 4:3) placed at a distance of about 57 cm from the chinrest. The height of the chin-rest, chair and monitor were adjusted in such a way that the center of the monitor and eyes lie on a horizontal plane. We sampled gaze location and pupil area using a video-based desktop-mounted infrared eye tracker at a frequency of 240 Hz (Model: ETL-200; ISCAN Inc., Woburn, MA, USA) that interfaced with TEMPO/VideoSync software (Reflective Computing, St. Louis, USA) in real time. The spatial resolution (root-mean square error) of the eye tracker was ~0.1° visual angle.

### Stimuli and task

A novel variant of ‘choice countermanding’ task (Middlebrooks and Schall, 2014; Indrajeet and Ray, 2019) was introduced to study the modulation in the pupil size during target selection and saccade planning when the execution of planned saccade was uncertain (Indrajeet et al., 2020). The schematic of the temporal sequence of events and behavior in the task is presented in Figure 1. Each trial began with the presentation of a fixation spot (~0.25° × 0.25° visual angle) of grey color inside a small square (~0.5° × 0.5° visual angle) of white color at the center of the monitor. The white square enclosing the fixation spot disappeared, and four checker boxes (~2.5° × 2.5° visual angle) appeared simultaneously at the periphery along with a broken white circle of radius ~1.8° visual angle around the central fixation spot, only if the participant maintained an uninterrupted gaze-fixation for a period of 200 – 500 ms. The circumferential gaps between adjacent segments of the broken circle were oriented to peripheral checker boxes; each gap subtended 0.7 radian of arc at the center. A total of 64 small squares of same size were arranged in eight rows and eight columns, and each square was painted in either cyan or magenta color to form a checker box. Each of these checker boxes was π/2 radian apart from any of its immediate adjacent checker boxes on an imaginary circle of radius of 12° visual angle with the origin at the center of the display monitor. To manipulate the difficulty in target selection, the proportions of magenta color across four checker boxes were distributed either from 80% to 20% at an interval of 20%, or from 70% to 40% at an interval of 10%. No particular spatial pattern was followed for color distributions; small squares painted in cyan and magenta color were placed randomly in each checker box in every trial. The central white square that would surround the fixation spot reappeared at the end of trial and remained on the screen to minimize dilation of pupil in dark during the interval of about 2 seconds between consecutive trials.

**Figure 1.**
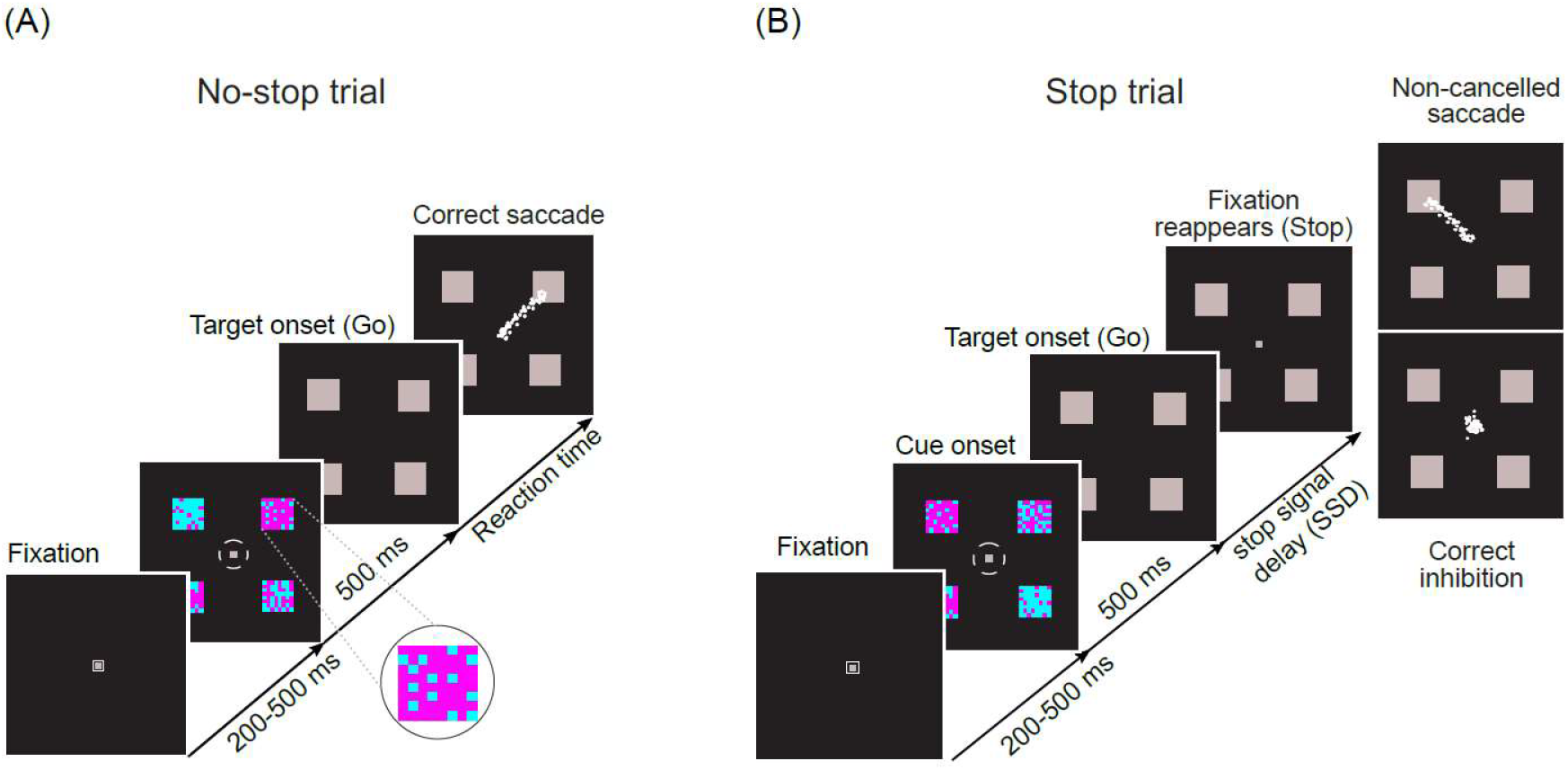
Schematic of temporal sequence of stimuli in a novel choice-countermanding task. Following a fixation period, four checker boxes painted with different proportion of cyan and magenta appeared peripherally along with a small white broken circle around the fixation spot. After 500 ms the fixation spot and the broken circle disappeared, and all four peripheral checker boxes were masked by grey squares simultaneously. (A) In the majority (60%) of trials (No-stop condition), participants were asked to select the checker box with the largest proportion of magenta color (magnified in inset), and orient their gaze to the location of the selected target following the disappearance of the fixation spot within a predetermined fixed period. Eye traces from representative saccades in no-stop trials are shown by white dots. (B) In the remaining trials (Stop condition), the fixation spot reappeared after a random delay (i.e. SSD) to instruct participants to withhold their eye-movement. In some trials, participants were able to inhibit pre-planned saccade successfully, but often they failed to do so resulting in the elicitation of non-cancelled saccades. No-stop and stop trials were randomly interleaved.

Participants were asked to assess carefully the proportion of magenta color in each of the checker boxes while maintaining their gaze at the fixation spot, and elicit saccadic eye movement to indicate their choice. Since PLR is a slow process with latency of 200 – 500 ms (Ellis, 1981), compared to saccade planning, we designed the task in a way that the pupillary muscles have enough time to respond to the visual decision cue before gaze orientation. After 500 ms of viewing, all checker boxes were masked by grey squares of size same as the size of colored boxes. The broken circle and the fixation spot both disappeared at the same time. The disappearance of the fixation spot served as a *go-signal* for the elicitation of a saccade towards one of the grey squares (place holders), which was at the location previously occupied by a checker box containing the maximum proportion of magenta squares. The disappearance of the broken circle at the central vision minimized possible arousal induced by the peripheral visual transient. In each trial, the center of the target was randomly selected from a set of 8 evenly distributed locations on an imaginary circle of 12° eccentricity, starting from 0 radian with respect to the horizontal meridian of the screen. All of the four grey squares remained on the screen until the trial ended. All stimuli were presented on black background. The luminance of magenta, cyan, and grey (mask) color stimuli used in the task was respectively 3.588 cd/m^2^ (CIE: x = 0.317, y = 0.169), 3.844 cd/m^2^ (CIE: x = 0.186, y = 0.239) and 3.541cd/m^2^ (CIE: x = 0.315, y = 0.343), and that of the black background was 0.245 cd/m^2^ (CIE: x = 0.312, y = 0.302).

In 60 % of total ~400 trials, referred to as *‘no-stop trials’,* participants were instructed to look at the selected peripheral target following the ‘go’ signal (i.e. the disappearance of fixation spot) as shown in Figure 1A. In the rest of the trials, referred to as *‘stop trials’* the fixation spot reappeared after a variable delay. The reappearance of the fixation spot acted as a ‘stop’ signal instructing the participants to refrain from looking at the peripheral target and keep their gaze fixed at the central fixation spot (Figure 1B). We refer to the delay between onset of the target and reappearance of the fixation spot as *‘stop signal delay’* or *SSD,* which varied from 100 to 600 ms at an interval of 100 ms with an infrequent jitter of about ±8 ms (i.e., half of the screen refresh duration at the refresh rate of 60 Hz). An interval of 2000 ms was introduced between consecutive trials.

In order to estimate a baseline reaction time, at the beginning of the experiment each participant was provided with 20 – 25 trials, wherein they performed only no-stop trials. While performing these trials the participants were unaware of the stop signal in the forthcoming primary recording session. This gave us an opportunity to estimate their natural saccadic reaction time (RT). After ~15 correct trials, the mean saccadic reaction time was calculated online, which was used as the baseline RT for the individual throughout the main recording session. Participants were not explicitly instructed to orient their gaze to the target as quickly as possible. Nevertheless, to discourage them from waiting for the stopsignal, we pseudo-randomly interleaved no-stop and stop trials. The deadline of the elicitation of saccade following the target onset was roughly 1.5 times the baseline RT rounded off to multiples of 100 ms, which ranged from 400 to 700 ms. Trials were scored online as correct if the participants looked at the peripheral target within the stipulated period in no-stop trials, or maintained the gaze on the central fixation spot for at least 300 ms after the stop-signal onset. Participants received feedback on their performance through an auditory tone of 1048 Hz for 200 ms in each correct trial. Total number of trials performed by an individual varied between 400 and 415. The amount of monetary rewards given to each participant was contingent on the total number of no-stop and stop trials performed correctly.

### Procedure

Programs written in Protocol Control Language (PCL) of TEMPO/ VideoSYNC software displayed the stimuli, sampled and stored the gaze location, pupil area and other task contingencies in real time, and provided auditory feedback at the end of each correct trial. A virtual square electronic window (~ 4° × 4°) around the central fixation spot specified the gaze-fixation region and another larger window (~ 5° × 5°) around the target specified the saccade-target region. A virtual electronic window around the central stop-signal was of the same size as the one around the saccade target. All offline analyses were performed using in-house programs written in Matlab® (The Mathworks Inc., USA). All statistical calculations were performed by using either Matlab® Statistical Toolbox (The Mathworks Inc., USA) or SigmaStat (SYSTAT Inc.,USA).

A boxcar window filter of length five was applied to smoothen the horizontal and vertical components of the eye positions. Subsequently, the program demarcated the onset of saccade offline when the eye-velocity and acceleration increased above 30°/s and 300°/s^2^ respectively, and the end of saccade was demarcated when the eye-velocity and deceleration decreased below the corresponding criteria for the beginning of saccade (Indrajeet and Ray, 2020). Trials with blink-perturbed saccades were excluded from subsequent analyses. The demarcations at the beginning and end of saccade in each valid trial were further scrutinized by visual inspection.

### ROC Analysis

To characterize the magnitude and time course of discrimination between pupil area in correct no-stop trials and correct stop trials, we used receiver operating characteristic (ROC) analysis based on signal detection theory (Green and Swets, 1966). The ROC curves were generated from two distributions of pupil area when saccades landed on the target and when planned saccades were cancelled. We randomly selected a unique set of 50 trials from both sets of trials to create ROC curves, and iterated until most of the trials in one set were exhausted. Comparisons were conducted by calculating ROC curves for successive bins of width 4 ms starting at 500 ms before the target onset and continuing up to 500 ms after the target onset. At every time bin, the proportion of trials in no-stop condition (i.e. sensitivity or true positive rate) was plotted against the proportion of trials in stop condition (i.e. 1 – specificity or false positive rate) that exhibited normalized pupil area larger than a criterion. The criterion was increased from the minimum to maximum at an interval of 0.001 to complete a ROC curve at that time bin. The area under the ROC curve (AUC) provides a reliable measure of the separation between distributions, with 0.5 indicating complete overlap and ±1 indicating complete separation. What AUC value describes a good discrimination? Hosmer et. al. (2013) noted that “Unfortunately there is no ‘magic’ number, only general guidelines”. The authors suggested AUC less than 0.7 is unacceptable, and “... not much better than a coin toss” (see p. 177).

### Neuromechanical homeomorphic model of PLR

Stark and colleagues pioneered mathematical models of PLR based on servo control mechanism of pupil (Stark and Sherman, 1957; Sun et al., 1983; Krenz and Stark, 1985). Simulation of servo-analytical models elucidated how different elastic properties and tension characteristics of pupillary muscle plant influence PLR dynamics (Semmlow and Stark, 1971; Semmlow and Chen, 1977). Usui and Hirata (1995) extended these ideas and introduced an elaborated homeomorphic model of PLR. “Homeomorphic models are those whose elements correspond to the anatomical, physiological, biomechanical, and neural elements of the experimental system” (Peterson et al., 1988). In this model, the characteristics of elastic element (passive elastic force), viscous element (viscous resistance), and contractile element (active force) were similar to Hill’s three-element model (Hill, 1938) of muscle in both circular sphincter and radial dilator muscle, respectively (Figure 6). The model determined the non-linear interactions between the dynamic properties of sphincter and dilator muscle components as they received inputs from the parasympathetic and sympathetic division of the autonomic nervous system (ANS), respectively. The following general equations described the elasticity, viscosity and characteristic of tension generator of muscles.

Elasticity:

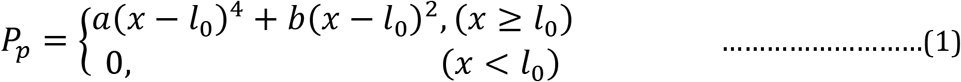

where *x* is the pupil radius, *a* and *b* are constants, and *l*_0_ is the length of muscle at the rest.

Viscosity:

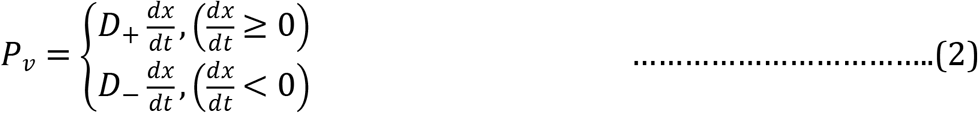

where D_+_ and D_−_ are the viscous coefficients at the phase of stretch and release, respectively.

Tension generator:

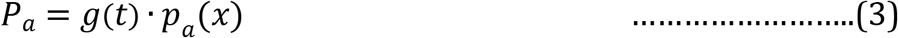

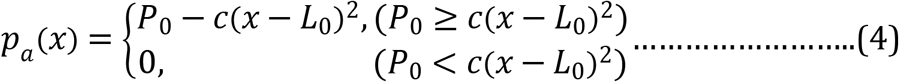

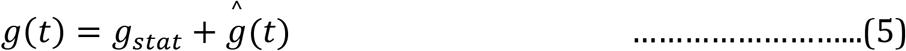

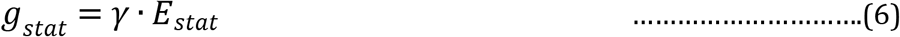

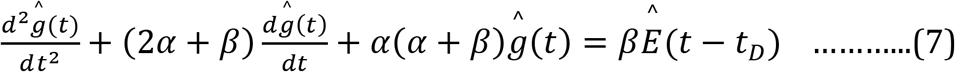

where *g*(*t*) and *P_a_*(*x*) are input dependent and muscle length dependent terms of active tension, respectively; *L_0_* is the muscle length at which maximum active tension *P_0_* is generated, respectively; *g_stat_* is the DC part of *g*(*t*), while *ĝ*(*t*) represents the AC part; similarly, E_stat_ is the DC part of autonomic activity *E*(*t*) and *Ê*(*t*) is the AC part; *t_D_* is the delay time of response; *C* and *γ* are constants; *α* and *β* are the time constants of the off and on slope of the isometric twitch response, respectively. Usui and Hirata (1995) simplified the two-dimensional plant structures into a onedimensional push-pull structure. A set of differential equations, as given below, was used to simulate the behavior of the pupil diameter to a flash of light. In these equations, subscript ‘s’ denotes sphincter parameters, subscript ‘d’ denotes dilator parameters, ‘stat’ and ‘^’ stand for static and dynamic part of the parameter; *x_max_* denotes the maximum radius of pupil. System equation of the model:

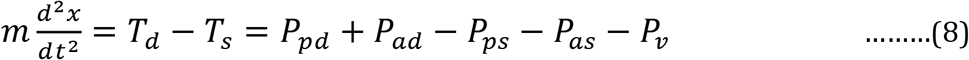

Dilator tension characteristics:

Passive tension

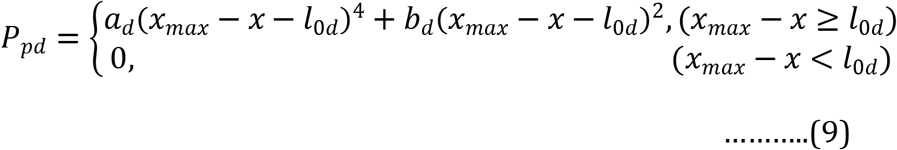

Active tension

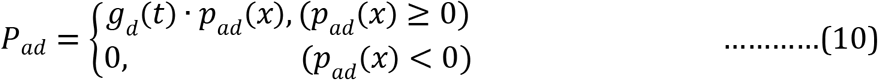

Elasticity

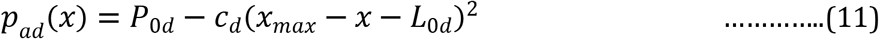

Dynamics

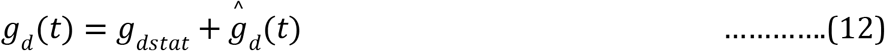

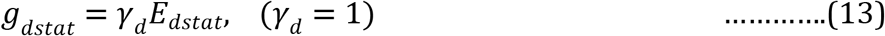

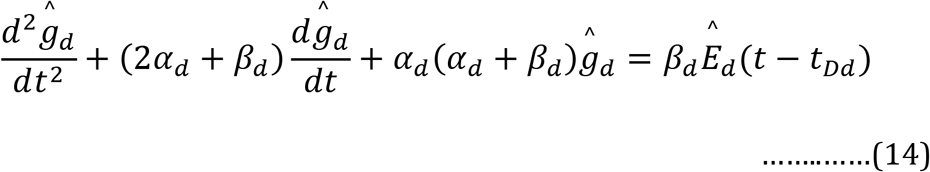

Sphincter tension characteristics:

Passive tension

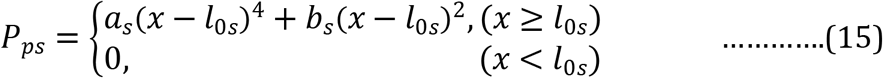

Active tension

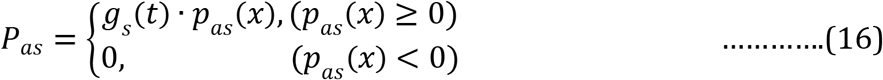

Elasticity

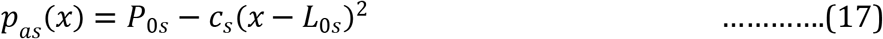

Dynamics

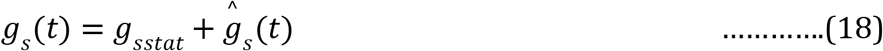

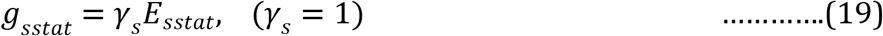

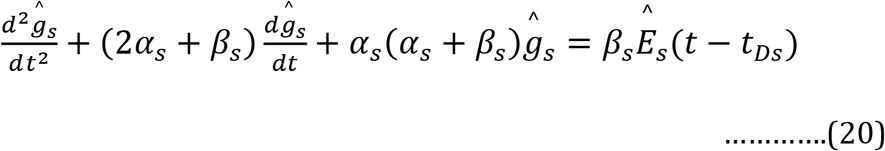

Total viscosity of the sphincter and dilator

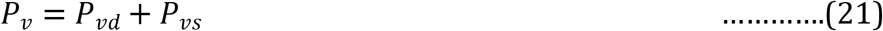

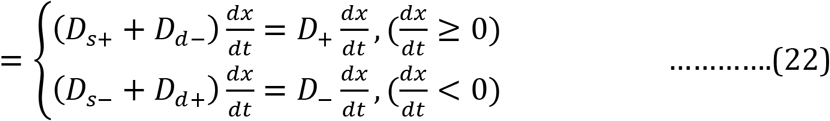

Autonomic Nervous innervation:

Sympathetic nervous activity

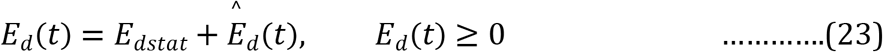

Parasympathetic nervous activity

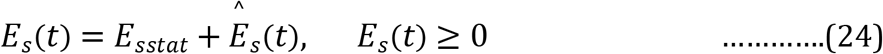

Reciprocal relationship between the parasympathetic and sympathetic nervous activities

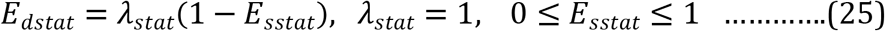

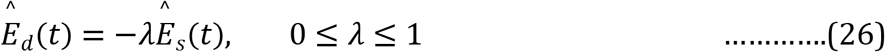

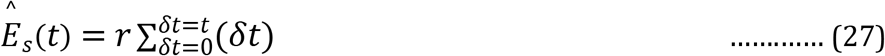

Studies on awake and behaving monkeys previously suggested a robust relationship between the saccade latency and the rapidness of the FEF and SC build up activity. The firing rate of neurons in these areas reaches a threshold before saccade onset (Hanes and Schall, 1996; Dorris et al., 1997). The saccade latency is inversely related to the rate with which the activity of movement (M) neurons arrives at a threshold. We assumed that the dynamic components *Ê_d_*(*t*) and *Ê_s_*(*t*) of sympathetic and parasympathetic division of ANS, as shown in equation 23 and 24, were driven by the rising activity 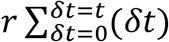 in the FEF and SC, respectively, where *r* was the rate of increase in activity (i.e. readiness of saccade derived from the reciprocal of saccade latency).

## RESULTS

### Performance and reaction time

In a decision making task, wherein participants were required to indicate their choice by orienting gaze, and refrain from elicitation of saccade in response to infrequent stop-signal (Figure 1), we considered correct no-stop and non-cancelled stop trials to study PLR dynamics. Saccade latency or reaction time (RT) in each trial was calculated by subtracting the time of the target (or go-signal) onset from the time of saccade onset. The appearance of the visual decision-cue that evoked PLR preceded the go-signal onset by 500 ms. The average (± SEM) percentage of correct no-stop and cancelled stop trials was 97.25 (± 0.41) and 47.1 (± 2.71), respectively (Figure 2A). The average (± SEM) overall saccadic reaction time (RT) in correct nostop and non-cancelled stop trials was 338 (± 12) ms and 344 (± 8) ms, respectively (Figure 2B). A paired *t-test* indicated no significant (P = 0.718) difference between the mean RT in correct no-stop trials and non-cancelled stop trials. Note that participants were allowed to initiate saccade only after the go-signal, which prevented them from eliciting very fast saccades that could largely escape inhibition in stop-trials. Given that PLR is a slow process relative to saccade planning, the introduction of go-signal was warranted. Therefore, having no significant difference between the mean saccade latency in rewarded no-stop and unrewarded stop trials was not surprising. In Figure 2C, the average (± SEM) percentage of error in stopping across the participants is plotted against stop-signal delay (SSD). A repeated measure one way ANOVA indicated a steady increase in error with increasing SSD. The main effect of SSD on the percentage of non-cancelled stop trials was found significant [F (5,95) = 74.153, P < 0.001]. Post-hoc pairwise multiple comparisons (Holm-Sidak method) revealed that the percentage of non-cancelled stop-trials in all possible pairs of SSD were significantly different from each other (minimum P < 0.001, maximum P = 0.023), except between the longest SSDs. In Figure 2D, the average (± SEM) RT in non-cancelled or error stop trials across the participants is plotted against stop-signal delay (SSD). A repeated measure one way ANOVA indicated increase in non-cancelled RT with increasing SSD, with a significant main effect of SSD on non-cancelled stop RT [F (5,92) = 17.205, P < 0.001]. Further, post-hoc pairwise multiple comparisons (Holm-Sidak method) showed that the average (± SEM) non-cancelled stop RT in all possible pairs of SSD were significantly different from each other (minimum P < 0.001, maximum P = 0.009), except between all adjacent pairs of SSD. An increase in percentage error and non-cancelled RT with increasing SSD together indicate that the participants correctly followed the instructions, and deliberately attempted to refrain from eliciting pre-planned saccade in stop trials.

**Figure 2.**
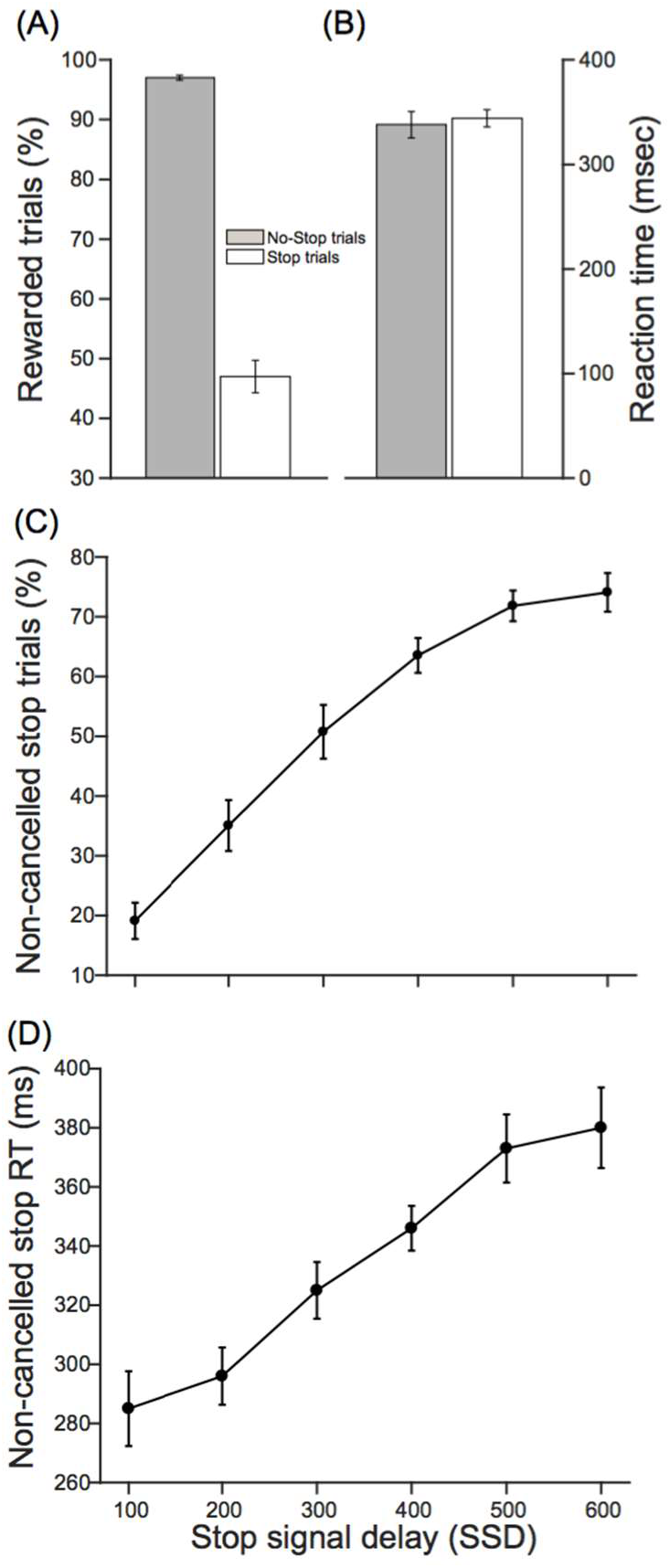
Performance and reaction time (RT) in correct no-stop and non-cancelled stop trials. (A) – (B) The average percentage of rewarded trials and reaction time (RT) in nostop (grey) and stop (white) condition. (C) – (D) The average percentage of failure in stopping saccade and reaction time in non-cancelled stop trials gradually increased as stop-signal delay (SSD) increased. Error bars indicate standard error of corresponding mean. These results ensure that the participants followed the instructions, and attempted to cancel planned saccade in stop trials.

### Pupillometry

Pupil size of participants was recorded in ISCAN eye-tracker’s arbitrary unit. In some trials, we observed discontinuity in the data wherein participants presumably blinked while making a choice and preparing for orienting gaze. In addition, in some trials, pupil size exhibited unusually extreme modulation and the difference between the maximum and the minimum exceeded 20×10^3^ in arbitrary unit. Those trials were removed from subsequent analyses because an abrupt modulation in pupil size might be task-irrelevant, for example, due to partial occlusion of pupil (Kret and Sjak-Shie, 2019). We normalized raw pupil area with respect to the average pupil area over a period of 100 ms that spanned from 200 ms to 100 ms before the cue onset. Subsequently, we aligned normalized pupil size either at the cue or saccade onset, smoothened and corrected the baseline to 1 (Mathôt et al., 2018). Finally, we performed a discrete Fourier transformation on pupil data in each trial to generate a power spectrum of the signal. As expected, the maximum power came from 0 Hz frequency because of the non-oscillatory nature of PLR dynamics. Note that the offset power is proportional to the square of the non-oscillatory component of amplitude of a time-varying signal. We removed outliers using conventional interquartile range (IQR) method. In total, we pooled 2776 trials (1900 no-stop trials and 876 non-cancelled stop trials) to understand the relationship between PLR dynamics and saccade planning.

We examined whether the pupil dynamics varies with saccadic reaction time. To this end, we first segregated all trials in which saccades were elicited into three subsets of nearly equal size with increasing RT (i.e., 33 and 66 percentiles of RT distribution were used to slice the distribution into three parts). The average (± SEM) RT measured relative to the target onset (i.e. 500 ms after cue onset) in short, medium, and long RT groups was 231 (± 2) ms, 320 (± 1) ms, and 453 (± 3) ms, respectively. Figure 3A shows differential modulation in the average (± SEM) normalized pupil size across trials in each of these RT groups for a second following the cue onset. To calculate the onset time of pupil constriction in each trial we used a sliding window of 100 ms (i.e. 25 data samples). Wilcoxon signed rank test was performed on pupil size within the time window, which was shifted rightward on the time axis by 4 ms (i.e. one data sample), in every iteration starting from the time of cue onset. We demarcated the onset of pupil constriction when the median pupil area significantly (P<0.05) decreased below the baseline 1, and continued to dwindle steadily for at least another 100 ms. Note that the slope me We also calculated the duration of pupil constriction in each trial by subtracting the time of constriction onset from the time when pupil size decreased to a minimum. Kruskal-Wallis one-way ANOVA on ranks suggested that the differences in the median values of constriction duration (i.e. 456, 456, and 444 ms) among trials in short, medium and long RT group were not significant (P = 0.601). The linear rate of pupil constriction in each trial was calculated by dividing the net change in pupil area by 300 ms interval from the constriction onset. The average (± SEM) onset time of pupil constriction relative to the cue onset in short, medium, and long RT groups was 309 (± 3) ms, 319 (± 3) ms, and 321 (± 3) ms, respectively. Therefore, addition of 300 ms to the average onset time of pupil constriction in each RT group was still less than the average onset time of saccade in that group relative to the cue onset. However, longer average saccade onset time does not necessarily ensure the gaze fixation was maintained during the period for which the rate of pupil constriction was calculated in every trial. In order to understand the pupil dynamics strictly before gaze orientation we repeated this analysis later aligning pupil size in each trial at saccade onset (Figure 4).

**Figure 3:**
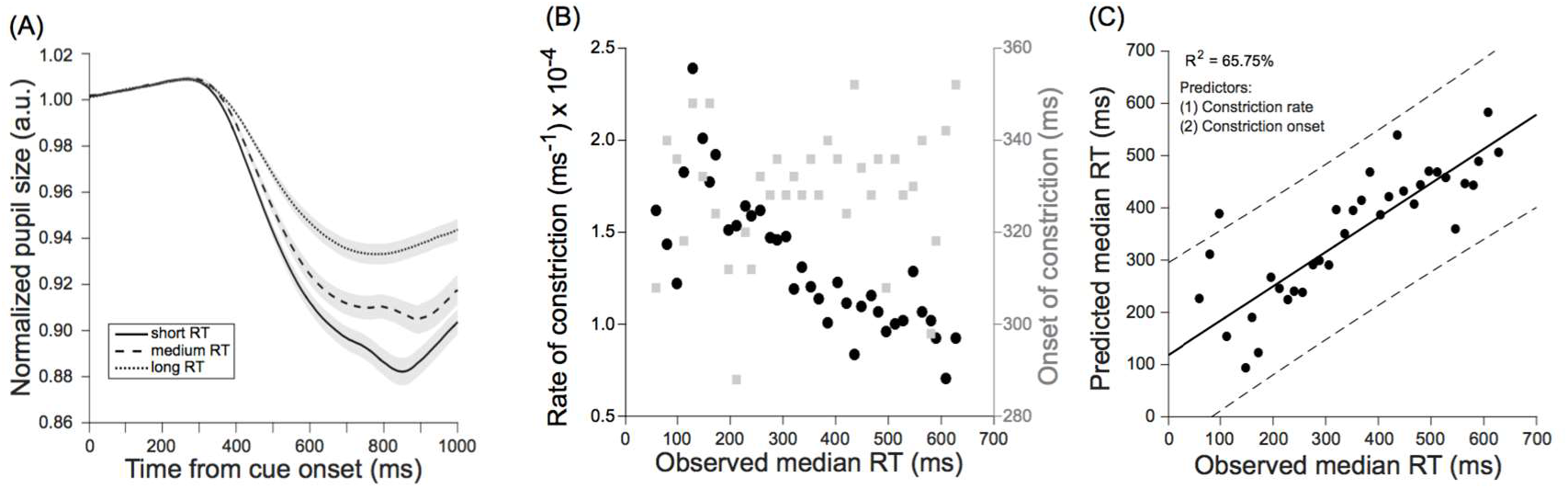
Influence of saccadic reaction time (RT) on pupil dynamics from the decisioncue onset. The average smoothened, normalized and baseline corrected pupil size in correct no-stop and non-cancelled stop trials together across the population (N=20) that yielded short (solid), medium (dashed) and long (dotted) RT exhibited differential dynamics. The grey patches are overlaid on the traces to show corresponding standard error of mean pupil size. (B) Median onset time (grey square) and rate (black circle) of pupil constriction are plotted against median RT of trials grouped into RT bins of 16 ms. The rate of pupil constriction was inversely related to RT of impending saccade. However, no such relationship between the onset time of pupil constriction and RT was found. (C) A linear model of RT with onset time and rate of pupil constriction as predictors accounted for 65.75% variance in the observed data. Dotted lines show 95% confidence interval of the best fit of predicted RT plotted against observed RT.

**Figure 4:**
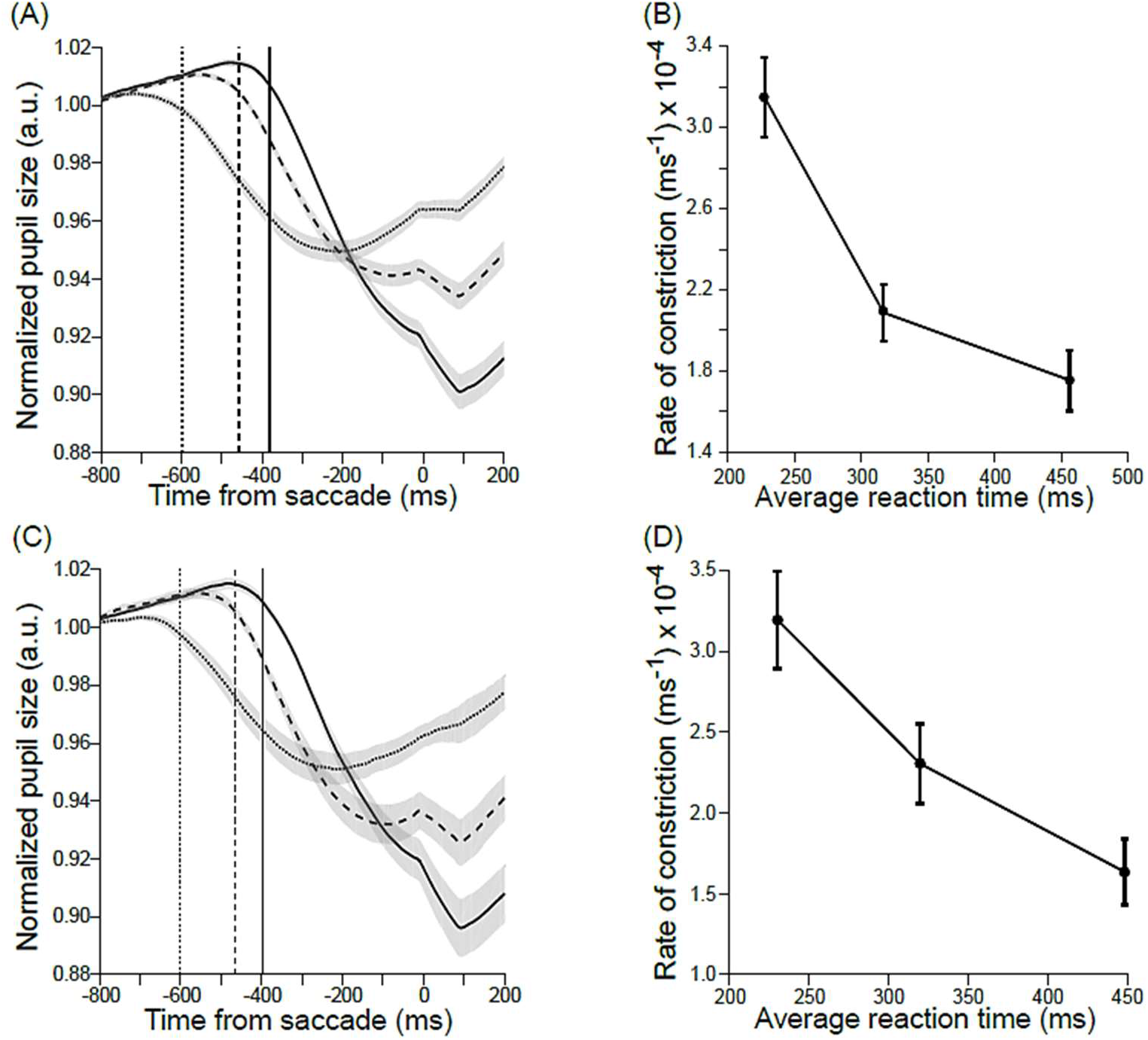
Influence of saccadic reaction time (RT) on pupil dynamics relative to saccade onset. The average smoothened, normalized and baseline corrected pupil size in (A) correct no-stop and (B) non-cancelled stop trials across the population (N=20) that yielded short (solid), medium (dashed) and long (dotted) RT. The grey patches are overlaid on the traces to show corresponding standard error of mean pupil size. The average (± SEM) linear rate of pupil constriction was calculated in 300 ms following the onset of pupil constriction across (C) correct no-stop and (D) non-cancelled stop trials in three RT groups. The average onset time of pupil constriction are shown by thin vertical lines (short RT: solid, medium RT: dashed, and long RT: dotted). The result shows that the pupils constricted at a higher rate during saccade planning when saccade elicitation was faster.

We sought to know whether the time of pupil constriction onset, or the rate of pupil constriction, or both, as shown in Figure 3A, can predict saccadic RT. In order to reduce the variability in the data, we binned trials that yielded RT within 16 ms (a refresh duration of 60 Hz display), and calculated median RT, median pupil constriction onset, and median pupil constriction rate in each bin (Figure 3B). A linear model of the form

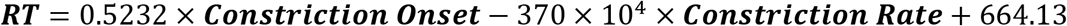

explained maximum 65.7% (R^2^ = 0.657, adjusted R^2^ = 0.637) variability in the observed median RT, which was statistically significant (P<< 0.001; F-statistic vs. constant model: 31.7). Significant contribution in predicting median RT came from the median rate of pupil constriction (P<< 0.001), but not from the median onset of pupil constriction (P = 0.669). This was further ascertained by Pearson’s coefficient (ρ) of correlation, which was significant between the median RT and the median constriction rate (ρ = –0.81, P<0.0001), but not significant between the median RT and the median constriction onset (ρ = 0.13, P = 0.447). Figure 3C contrasts predicted and observed median RTs.

We further tested whether the inverse relationship between saccade latency and pupil constriction rate remained consistent irrespective of the cognitive context. We segregated trials in each of two cardinal sets – correct no-stop trials and non-cancelled stop trials, into three subsets of trials of nearly equal size, with increasing RT. In order to understand the pupil dynamics strictly before gaze orientation, the average (± SEM) pupil area in different RT groups of correct no-stop trials and noncancelled stop trials aligned on saccade onset are plotted in Figure 4A and 4C, respectively, and corresponding mean onset time of pupil constriction for each of these groups is shown by a thin vertical line. The onset and rate of pupil constriction in each trial was calculated following the procedures described above. In each RT group, the average (± SEM) rate of constriction in the pupil size was calculated and plotted against corresponding average RT for correct no-stop trials and noncancelled stop trials in Figure 4B and 4D respectively. The median rate of pupil constriction for short, medium and long RT group was 0.139, 0.106 and 0.093 arbitrary unit per second respectively for correct no-stop trials, and 0.125, 0.108 and 0.095 arbitrary unit per second respectively for non-cancelled stop trials. Kruskal-Wallis one way ANOVA on rank was performed on the rate of pupil constriction. The median values of the rate of pupil constriction in three RT groups of trials in both correct no-stop [H(2) = 69.27] and non-cancelled stop [H(2) = 34.84] conditions were significantly different (P<0.001). A post-hoc multiple comparisons using Dunn’s method suggested that the pupil constriction rate in long, medium and short RT groups in both correct no-stop trials and non-cancelled stop trials were significantly (P < 0.05) different from each other. Our data thus clearly indicate that the rate of pupil constriction inversely varied with the latency of impending saccade regardless of the reward outcome.

### Attentional orientation or motor preparation?

In order to resolve the issue whether observed modulation in pupil dynamics with varying saccade latency was due to covert shift of attention (Mathôt et al., 2013) or motor preparation (Mathôt et al., 2015), we compared pupil size aligned at the target onset in correct no-stop trials wherein saccades were elicited with that in correct stop trials wherein saccades were cancelled (Figure 5A). Earlier studies suggested that obligatory pre-saccadic shift of attention occurs if and only if saccades are inevitable (Ray et al., 2009; Born et al., 2014). Therefore any reasonable difference in the pupil dynamics in two sets of trials would reflect influence of covert attentional shift on pupil size. We performed receiver operating characteristic (ROC) analysis and computed area under the ROC curve or AUC, which never reached to a desired level of discrimination (0.7) as shown in Figure 5B. Note that, discrimination with 0.5 < AUC ≤ 0.7 is anecdotal and result should be interpreted with caution (Hosmer Jr et al., 2013). Therefore it appears unlikely that shift of attention contributed to modulation of visually evoked PLR, instead saccade preparation may be the key contributor to this modulation.

**Figure 5:**
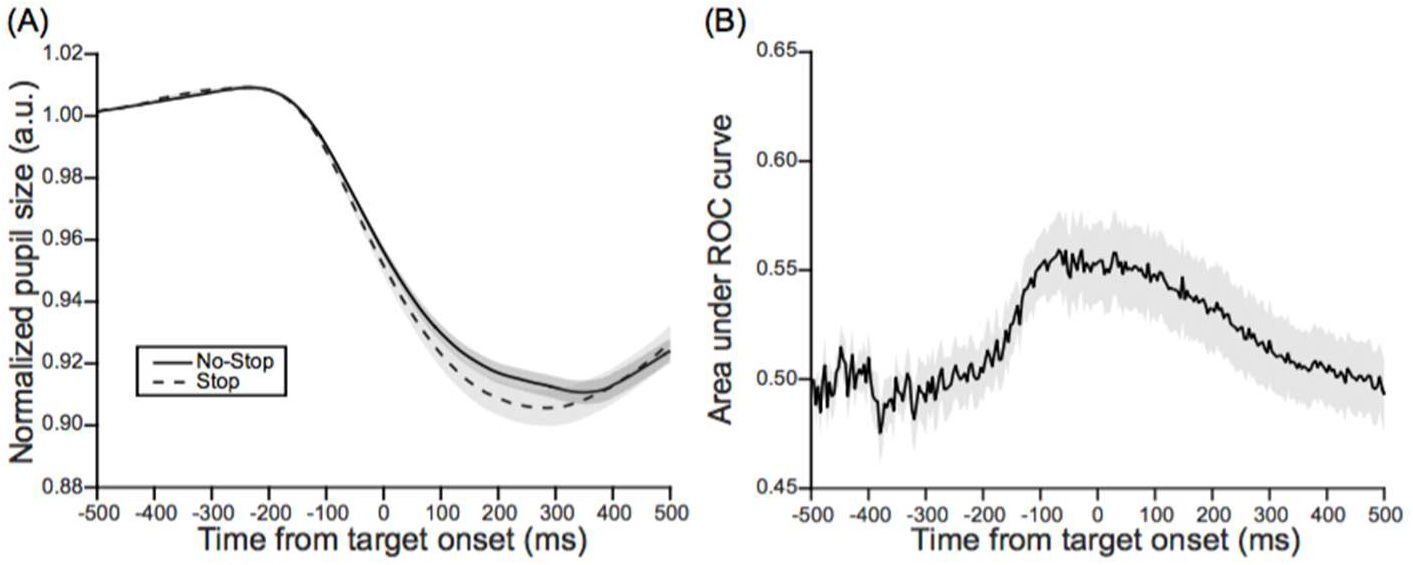
(A) Pupil dynamics in correct no-stop (solid line) and stop (broken line) trials aligned at target onset. (B) Corresponding area under the ROC curve *(see* MATERIALS AND METHODS). Grey patches indicate standard error of mean.

**Figure 6:**
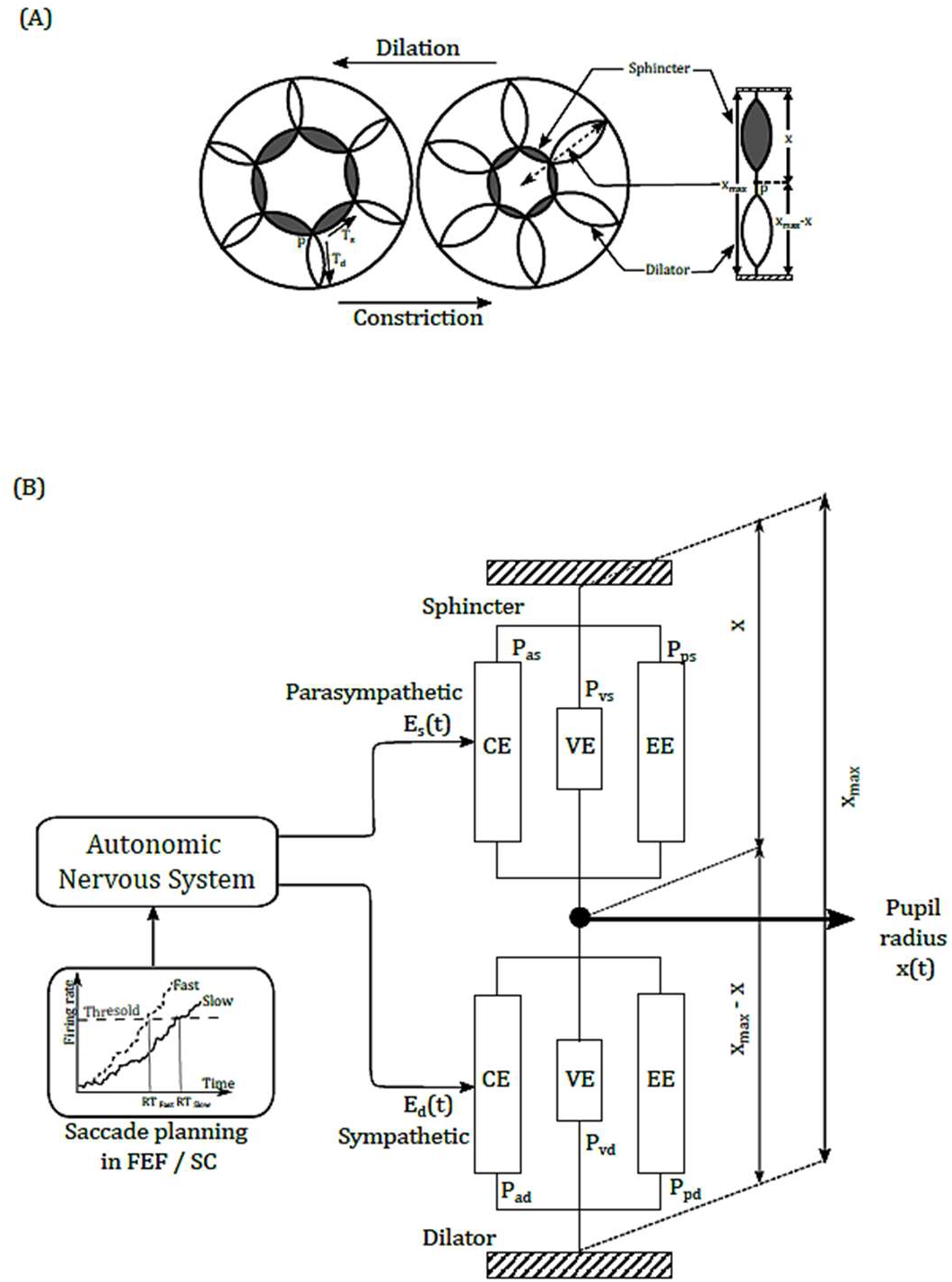
(A) Schematic of radial arrangement of pupillary muscles shown by shapes similar to loom spindle (sphincter: grey, dilator: white) in dilated (left) and constricted (middle) condition of pupil aperture (inner circle), and a one-dimensional mechanical equivalent of two-dimensional pupillary muscles (right). (B) Schematic of revised Usui-Hirata (1995) model of pupillary muscle plant. In this model, rise-to-threshold activity in the FEF and SC (shown in inset) during saccade planning determined the dynamic part of parasympathetic (E_s_) and sympathetic (E_d_) autonomic activity that constricts and dilates pupil, respectively. T_s_ = tension in sphincter; T_d_ = tension in dilator; p = T_d_ – T_s_; x = radius of pupil aperture; xmax = total length of sphincter and dilator; CE = contractile element; VE = viscous element; EE = elastic element; P_as_ = active tension in sphincter, P_ps_ = passive tension in sphincter, P_vs_ = viscosity in sphincter; P_ad_ = active tension in dilator, P_pd_ = passive tension in dilator, P_vd_ = viscosity in dilator; E_s_(t) = time dependent parasympathetic activity; E_d_(t) = time dependent sympathetic activity.

### Model simulation

We revised and simulated Usui-Hirata (1995) model of pupillary muscle plants (Figure 6). In this revised model, activity in the FEF and SC building up over time at a steady rate controlled the dynamic component of sympathetic *Ê_d_*) and parasympathetic (*Ê_s_*) activity (see equation 23 and 24), which in turn controlled the dynamics of tension generated in the dilator and sphincter muscle plant in the iris (see equation 14 and 20), respectively (see MATERIALS AND METHODS for more information on the model). It is important to note that in the model, activity in the FEF/SC at any given instant, not the rate of activity buildup, determined the dynamics of autonomic drive (i.e. *Ê_s_* and *Ê_d_*) to the pupillary muscle plant. Therefore the rate of change in simulated pupillary aperture should not be mistaken as the rate of saccade planning. We simulated the model using Matlab Simulink® 8.6 (The Mathworks Inc., USA) software running on an iMac (Apple Inc., USA) computer with 3.2 GHz Intel® Core i5 processor, 8 GB RAM, and OSX 10.11.4 operating system. Simulation of the model continued for 1500 ms at an equal interval of 4 ms.

In each trial from a set of 600 trials, the build up activity in the FEF and SC was 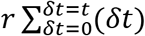, where the rate of accumulation *r* (i.e. readiness of saccade) was sampled from a normal distribution. The minimum and maximum value of *r* was determined by the reciprocal of maximum (800 ms) and minimum (50 ms) allowed saccadic reaction time respectively. We considered that the visual delay in the FEF was 100 ms (Pouget et al., 2005). The motor decision process in the FEF could evolve anytime during the remaining 400 ms of exposure to the decision-cue. Therefore we assumed that the saccade related build-up activity in the FEF began a total of 300 ms (i.e. 100 ms visual offset plus on average 200 ms decision process offset) after the cue onset, which we referred to as visuo-movement offset (a constant VM) in the model. Table 1 shows values of other parameters of the model. Note that homeomorphic models “typically have more parameters than the less realistic phenomenological or input/output models” (Peterson et al., 1988)

**Table 1:** Parameters of the neuro-mechanical model of pupillary muscles. Description of these parameters and value used for simulation are enlisted. Symbols corresponding to these parameters in the equations used to design the model are given in ‘Materials and Methods’. Parameters shown in shaded rows are either modified or introduced to incorporate influence of saccade planning on autonomic input to pupillary muscle plants in Usui-Hirata (1995) model of PLR.

| *Model parameter* | *Value* | *Description* | *Referring equation* |
| --- | --- | --- | --- |
| $\alpha_s$ | 3.50 | Off slope of isometric twitch of sphincter | 20 |
| $\alpha_d$ | 0.50 | Off slope of isometric twitch of dilator | 14 |
| $\beta_s$ | 8.00 | On slope of isometric twitch of sphincter | 20 |
| $\beta_d$ | 1.50 | On slope of isometric twitch of dilator | 14 |
| $t_{Ds}$ | 0.01 | Visual delay to parasympathetic PLR pathway | 20 |
| $t_{Dd}$ | 0.01 | Visual delay to sympathetic PLR pathway | 14 |
| $a_s$ | 0.10 | Passive tension coefficient for sphincter | 15 |
| $a_d$ | 0.70 | Passive tension coefficient for dilator | 9 |
| $b_s$ | 0.35 | Passive tension coefficient for sphincter | 15 |
| $b_d$ | 0.75 | Passive tension coefficient for dilator | 9 |
| $c_s$ | 100.00 | Elasticity coefficient for sphincter | 17 |
| $c_d$ | 15.00 | Elasticity coefficient for dilator | 11 |
| $l_{0s}$ | 1.50 | Length of sphincter at rest | 15 |
| $l_{0d}$ | 1.50 | Length of dilator at rest | 9 |
| $L_{0s}$ | 3.50 | Length of sphincter at which $P_{0s}$ is generated | 17 |
| $L_{0d}$ | 3.50 | Length of sphincter at which $P_{0d}$ is generated | 11 |
| $x$ | 2.175 | Initial radius of pupil | 1, 4, 9, 11, 15, 17 |
| $x_{max}$ | 4.00 | Maximum radius of pupil | 9, 11 |
| $P_{0s}$ | 400.00 | Maximum active tension in sphincter | 17 |
| $P_{0d}$ | 120.00 | Maximum active tension in dilator | 11 |
| $D_+$ | 14.00 | Viscous coefficients at the phase of stretch | 2, 22 |
| $D_-$ | 70.00 | Viscous coefficients at the phase of release | 2, 22 |
| $E_{sstat}$ | 0.15 | Static component of parasympathetic activity | 24, 25 |
| $E_{dstat}$ | 0.85 | Static component of sympathetic activity | 23, 25 |
| $\hat{E}_s(t)$ | Dynamic | $r \sum_{\delta t=0}^{\delta t=\tau}(\delta t)$, dynamic component of parasympathetic autonomic activity | 24, 26 |
| $\hat{E}_d(t)$ | Dynamic | $-r\lambda \sum_{\delta t=0}^{\delta t=\tau}(\delta t)$, dynamic component of sympathetic autonomic activity | 23, 26 |
| $\lambda$ | 1 | Gain | 26 |
| $r$ | Random | Readiness (1÷saccade latency) | 27 |

Figure 7A shows pupil dynamics (black lines) in three simulated trials for three different saccade latencies (100, 276, 436 ms) relative to the go-signal onset. Simulated build-up activity (grey lines) in the FEF and SC corresponding to these trials are overlaid. This illustrates how efficiently the speed of saccade planning modulated pupil size in the model. We sorted simulated trials based on simulated RT, and separated into three sets of 200 trials each in ascending order of the average (± SD) RT, which was 217(±63), 441(±68) and 673(±66) ms. Figure 7B shows the average (± SD) simulated normalized pupil size in these sets of trials from the beginning to next one second of simulation. The simulated data mimicked the primary empirical finding of this study that the rate of pupil constriction inversely varied with saccade RT (see Figure 3A). We ascertained this by calculating the rate of pupil constriction for the first 200 ms from the onset of pupil constriction in each trial. The median rate of pupil constriction for short, medium and long RT group was 0.256, 0.210 and 0.178 arbitrary unit per second respectively. Kruskal-Wallis one way ANOVA on rank was performed on the rate of pupil constriction. The median values of the rate of pupil constriction in three RT groups of trials were significantly different [H(2) = 532.45, P<0.001]. A post-hoc multiple comparisons using Tukey method suggested that the pupil constriction rate in long, medium and short RT groups were significantly (P < 0.05) different from each other. The average (± SD) rate of simulated pupil constriction in three RT groups of trials are plotted against corresponding average RT in Figure 7C. This result not only establishes a robust relationship between saccade planning and pupil size, but provides a neuromechanical account of the relationship, which is quite physiologically feasible.

**Figure 7:**
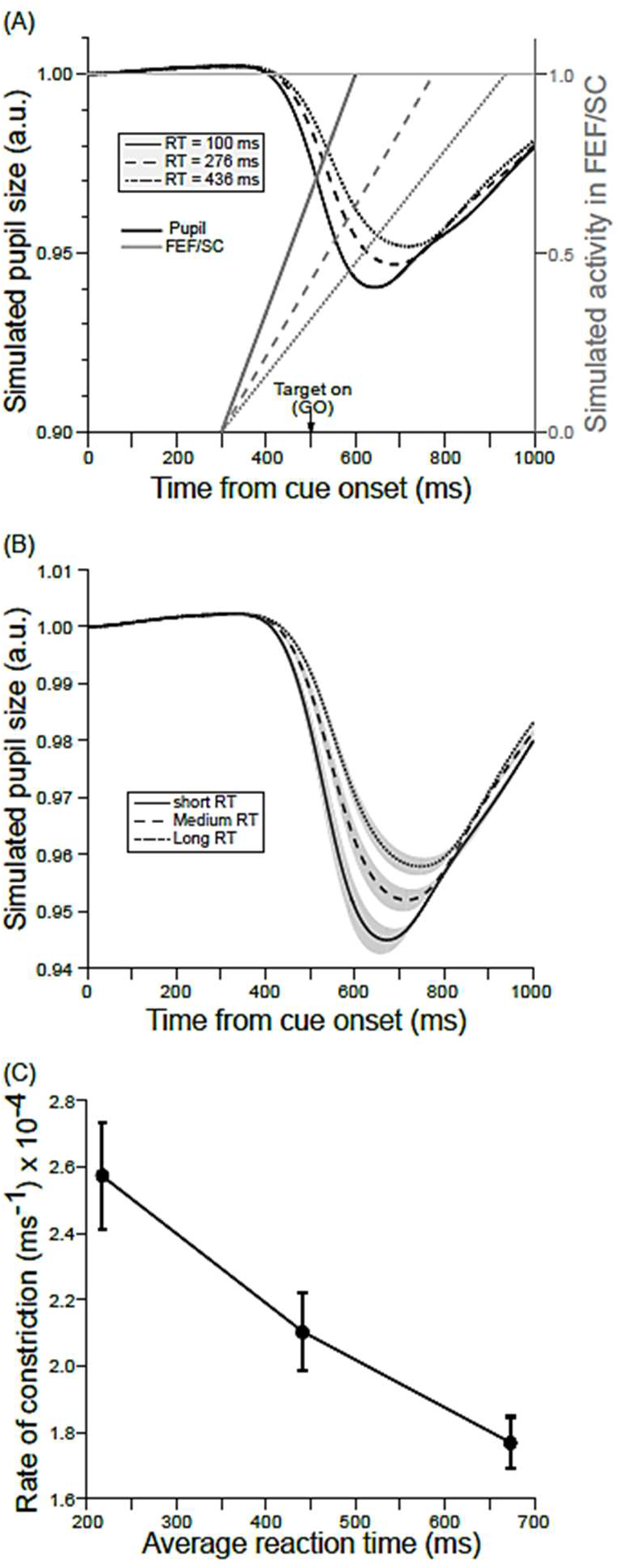
Influence of saccadic reaction time (RT) on simulated pupil dynamics from the decision-cue onset. (A) Simulated rise-to-threshold FEF/SC activity (grey) in three exemplar trials that yielded RT of 100, 276 and 436 ms are overlaid with corresponding simulated normalized and baseline-corrected pupil size (black) for short (solid), medium (dashed) and long (dotted) RT. A horizontal thin grey line at 1.0 indicates the threshold of saccade elicitation. RT was calculated relative to the target onset (down arrow). (B) The average (± SD) simulated normalized and baseline-corrected pupil size across trials that yielded short (solid), medium (dashed) and long (dotted) RT. (C) The average (± SD) rate of simulated pupil constriction calculated over 200 ms from onset of constriction in three RT groups are plotted. Simulation of the neuro-mechanical model of pupillary muscles mimics the behavioral data (see Figure 3A) and supports the primary empirical finding that the constriction rate is inversely related to RT.

## DISCUSSION

The pupils involuntarily and consensually constrict in light and dilate in dark. Interestingly, PLR is subjective; pupil size depends on perceived luminance instead of the actual one (Binda et al., 2013b). Similar to saccade, pupillary response to luminance is partly reflexive and partly under cognitive control (Einhäuser, 2017; Mathôt, 2018). The non-visual factors, for example, mental effort, memory load, attention, arousal, time-perception, decision-making, environmental regularities too affect pupil size (Hess and Polt, 1964; Kahneman and Beatty, 1966; Bradshaw, 1967; Wierda et al., 2012; Alnæs et al., 2014; Binda et al., 2014; de Gee et al., 2014; Suzuki et al., 2016; Schwiedrzik and Sudmann, 2020). The current study has addressed a fundamental question: how does an imminent voluntary eye-movement influence the dynamics of involuntary pupillary light reflex (PLR)? We found that the rate of pupil constriction triggered by visual stimulation (i.e. the appearance of the decision cue) was inversely proportional to the latency of impending saccade – the shorter the response time, the faster the constriction of the pupils, and vice versa. Note that the same luminosity across the trials triggered PLR. The relationship between the PLR dynamics and saccade latency was independent of the reward outcome of saccadic behavior. This finding suggests that the brain areas that plan eye movement and make connections to PLR pathways play a critical role in light adaptation to maintain a stable percept of the world of coherent luminance around the time of the gaze shift.

The PLR dynamics inversely resembled neural dynamics of the FEF/SC activity during saccade preparation, indicating a strong coupling between pupil size and saccade planning. Our study confirms what was speculated earlier; that is the pupil size possibly be an effective proxy of neural activity in the SC (Wang et al., 2015). The FEF and SC are two important structures in the oculomotor network of the primate brain, which directly participate in saccade planning (Schall et al., 2002; Basso and May, 2017) and link cognition to saccade (Funahashi, 2014; Matsumoto et al., 2018). Earlier studies have shown that, in the FEF and SC, the activity of movement neurons for correct and non-cancelled errant saccades in their response field are comparable (Hanes et al., 1998; Paré and Hanes, 2003). The current study also shows that the pupil dynamics in two task-conditions are similar.

In a recent study on perceptual decision-making, the average latency of a button press was greater when their baseline (i.e., before onset of the visual decision cue) pupil size was larger indicating higher tonic baseline arousal level. In contrast, higher phasic arousal indicated by task evoked pupil dilation of greater magnitude accompanied faster button press (van Kempen et al., 2019). Similarly, larger pupil size prior to the target appearance accompanied slightly longer prosaccade latencies (Cherng et al., 2020). Pupil size distinguished between pro- and anti-saccade, and pre-target pupil dilation of greater magnitude in each task-type accompanied saccades with shorter latency (Wang et al., 2015). Further study is required to test whether PLR indeed influences movement latency independent of the effector.

Pupil size is controlled by a balanced and synergistic interaction between parasympathetic (constriction) and sympathetic (dilation) division of the autonomic nervous system, which innervate sphincter pupillae and radial muscle of iris respectively (Kardon, 2011). A steady increase in the excitatory innervation to sphincter pupillae modulated by the oculomotor system likely is the reason behind such dynamics of PLR. This has been replicated in our simulation study. What might be the neural network mediating a link between the pupil size and build-up activity in the FEF during saccade planning? Our current understanding of the PLR network allows us to envisage a pathway that mediates the translation of the modulation in neural dynamics to pupil dynamics during saccade planning (Figure 8).

**Figure 8:**
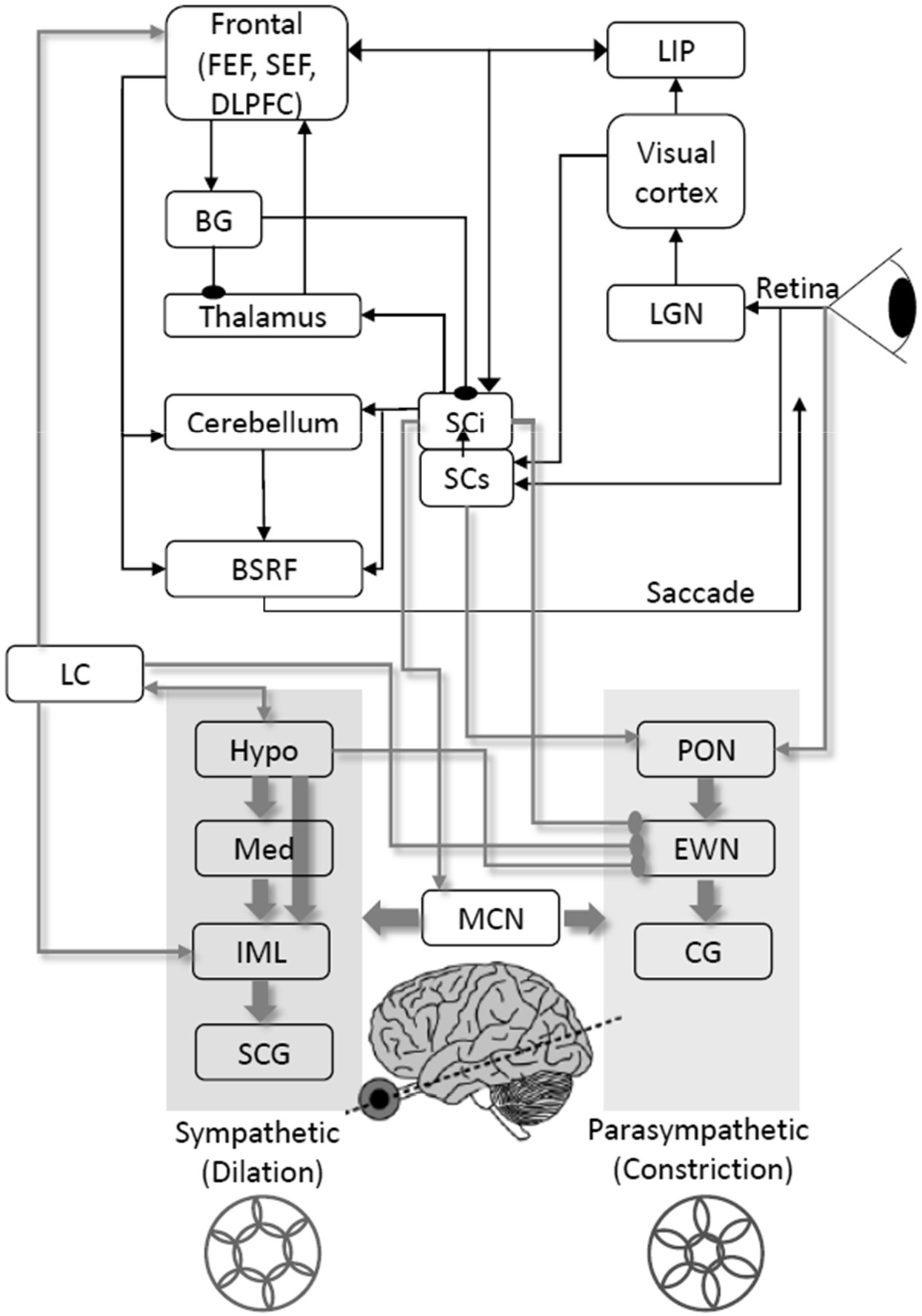
A plausible neural network for the realization of influence of planning saccadic eye movement on pupillary light response [based on (Munoz, 2002; Wang and Munoz, 2015; Mathôt, 2018)]. Abbreviations: frontal eye field (FEF), supplementary eye field (SEF), dorsolateral prefrontal cortex (DLPFC), lateral intra-parietal (LIP), basal ganglia (BG), lateral geniculate nucleus (LGN), superior colliculus intermediate (SCi), superior colliculus superficial (SCs), brainstem reticular formation (BSRF), locus coeruleus (LC), hypothalamus (Hypo), medulla (Med), intermediolateral nucleus of spinal cord (IML), superior cervical ganglia (SCG), pretectal olivary nucleus (PON), Edinger-Westphal nucleus (EWN), mesencephalic cuneiform nucleus (MCN), ciliary ganglion (CG). Arrows: excitatory (spear head), inhibitory (dot head), black (saccade pathway), gray shadowed (PLR pathway).

The FEF and SC in primate’s brain make necessary connection to modulate activity in the pretectal olivary nucleus (Harting et al., 1980; Leichnetz, 1982, 1990) that directly receives input from the retina (Güler et al., 2008), and projects to the Edinger–Westphal (EW) nucleus (Kourouyan and Horton, 1997). Preganglionic parasympathetic neurons in EW nucleus synapse with the ciliary ganglion that causes pupil constriction by innervating the iris sphincter muscle (Kardon, 2005). It appears that the SC_i_ receives projections from the FEF and excites the EW nucleus on the parasympathetic constriction pathway. Through efferent projections to the mesencephalic cuneiform nucleus (MCN), the SC_i_ also inhibits the EW nucleus and influences sympathetic dilation pathway. The pontine locus coeruleus (LC) may mediate tipping the balance between competing pathways through its efferent projection to the SC_i_ (Harting, 1977; Wang and Munoz, 2015). The moment-by-moment modulation in the baseline firing rate of LC has been implicated in the fluctuation of the pupil size (Aston-Jones and Cohen, 2005) and arousal (Breton-Provencher and Sur, 2019). Another possible network, although not clearly understood anatomically, involves the paragigantocellularis nucleus (PGi) of the ventral medulla that receives widespread inputs from cortical and subcortical areas and sends projections to the EW nucleus and influence dilation pathway via the LC (Gilzenrat et al., 2010; Joshi et al., 2016).

The norepinephrine or NE (also called noradrenaline) networks in primate’s brain are highly distributed having the LC as an area in it. The activity in the LC-NE pathway has direct positive moment-by-moment correlation with pupil aperture (Aston-Jones and Cohen, 2005). Dextroamphetamine and atomoxetine are used for the treatment of attention-deficit/hyperactivity disorder (ADHD) (Millichap, 2009). While dextroamphetamine causes the direct release of NE, atomoxetine is a selective reuptake inhibitor for NE (Kenny et al., 2015). Controlled application of d-amphetamine improves inhibitory control of both eye and manual movement in healthy humans (Allman et al., 2010). In a combined pharmacological-fMRI study it was found that the application of atomoxetine improved inhibitory control of human volunteers and increased activity of the right inferior frontal gyrus (rIFG) (Aston-Jones and Gold, 2009; Chamberlain et al., 2009). Therefore, the measurement of the pupil aperture as a proxy for the LC activity may reveal the progress in response inhibition. An ideal biomarker, which is an indicator of biological process or state, should be able to distinguish between the normal and disease state, be non-invasive and objective. Our findings indicate that the pupil size may index saccadic decision time, and may be used as a biomarker for disease with impairments in motor inhibition, for example, in ADHD, schizophrenia and Parkinson’s disease (Chamberlain et al., 2006; Joti et al., 2007; Wang et al., 2016; Thakkar et al., 2018). In short, taking aforementioned neurophysiological, psychopharmacological studies and our present study together, it appears that the FEF and SC strongly influence the pupil dynamics to optimize perisaccadic visual perception and minimize saccade transients (Mostofi et al., 2020).

## Authors’ contribution

PP collected and analysed data, and performed statistics. SR designed and programmed the task. SR designed and simulated the model. Both authors wrote the manuscript.

## Acknowledgements

This work was supported by a grant from the Wellcome Trust DBT India Alliance [IA/I/13/2/501015].

## Data and software accessibility

Data and the model designed in Matlab Simulink may be shared on request for legitimate purpose.

